# Recombinant single-cycle influenza virus as a new tool to augment antitumour immunity with immune checkpoint inhibitors

**DOI:** 10.1101/2021.07.31.454512

**Authors:** Matheswaran Kandasamy, Uzi Gileadi, Pramila Rijal, Tiong Kit Tan, Lian Ni Lee, Jili Chen, Gennaro Prota, Jing Zhang, Terence Rabbitts, Alain Townsend, Vincenzo Cerundolo

## Abstract

Virus-based tumour vaccines offer many advantages compared to other antigen delivering systems. They generate concerted innate and adaptive immune response, and robust CD8^+^T cell responses. We engineered a non-replicating pseudotyped influenza virus (S-FLU) to deliver the well-known cancer testis antigen, NY-ESO-1 (S-NY-ESO-1 FLU). Intranasal or intramuscular immunization of NY-ESO-1 S-FLU virus in mice elicited a strong NY-ESO-1 specific CD8+T cell response in lungs and spleen that resulted in the regression of NY-ESO-1 expressing lung tumour and subcutaneous tumour respectively. Combined administration with anti PD-1 antibody, NY-ESO-1 S-FLU virus augmented the tumour protection by reducing the tumour metastasis. We propose that the antigen delivery through S-FLU is highly efficient in inducing antigen specific CD8+T cell response and protection against tumour development in combination with PD-1 blockade.

## Introduction

The fight against cancer remains unfinished as it continues to be a major threat to human life. Tumour antigen specific strategies such as the dendritic cell (DC) vaccine^1^ have been previously investigated to elicit antigen specific anti-tumour immunity. Such approaches yielded limited success because of the profound tumour suppressive environment. Recent successes of treatments with immune checkpoint inhibition are impressive but leave many patients unaffected. Hence, multiple approaches are warranted to reverse tumour immunosuppression and induce anti-tumour T cell responses. The generation of antitumor immunity together with the reversal of tumour immune suppression might be achieved by triggering innate immune receptors which have been evolved to detect pathogen-associated molecular patterns (PAMPs).

Intrinsically immunogenic pathogens with potential pattern recognition receptor (PRR) agonistic functions can induce potent anti-tumour responses ^2^ and with the help of genetic engineering, the immunogenic pathogens can be exploited as vectors to deliver tumour associated antigens (TAA). Viruses are naturally immunogenic and represent an attractive vehicle for antigen delivery as several studies have shown that antigens expressed by virus are more immunogenic than soluble antigen administered with adjuvant^3, 4^. Moreover, viruses infect antigen-presenting cells (APCs) and express their transgenes^5–9^. The proinflammatory immune response induced by the virus infection enhanced the transgene or tumour antigen presentation which has led to not only an increase in the frequency of antigen-specific cytotoxic T cells but also their avidity to tumour antigen-expressing cancer cells^10^.

Influenza A virus (IAV) is an interesting candidate for antigen delivery since IAV infection elicits a strong antigen specific CTL response. We developed previously a pseudotyped replication deficient influenza A/Puerto Rico/8/34 (PR8) virus in which the HA signal sequence was inactivated (S-FLU)^11^; this virus can replicate only in the cell line expressing HA which provides the HA protein on pseudotyped virus particles for their binding to cells and importantly, the HA expressed by the cell line can be derived from any virus sub-type. Experimental infection with S-FLU virus has shown a robust CTL response but in the absence of neutralizing antibodies^11^. Here, we generated a recombinant S-FLU virus expressing NY-ESO-1(New York oesophageal squamous cell carcinoma 1) and evaluated its immunogenicity and therapeutic efficacy in preclinical mouse models. NY-ESO-1 is a well-known cancer-testis antigen (CTA). Its expression is normally restricted in germ cells, but it is highly dysregulated in some malignant cells ^12^. Given the immune-privileged nature of germline cells, NY-ESO-1 may be therapeutically targeted without substantial risk of immune related off-targeted effects. We found that intranasal or intramuscular immunization with NY-ESO-1 S-FLU virus elicits a robust NY-ESO-1 specific CTL response and suppresses the NY-ESO-1 expressing tumour development and spontaneous metastasis. Moreover, with anti PD1 antibody co-administration, NY-ESO-1 S-FLU virus immunization displays an enhanced tumour reduction.

## Results

### NY-ESO-1 S-FLU virus is able to express exogenous protein and induce a ^i^potent antigen specific CD8+T cell response

We generated S-FLU virus to express NY-ESO-1 by replacing the coding region of HA with NY-ESO-1 expressing sequence as described previously^11^(Fig.1A left). Three positive S-FLU clones were identified (Figure 1A (right panel)) and clone 2A9 was chosen and expanded for further experiments.

**Figure 1A.**
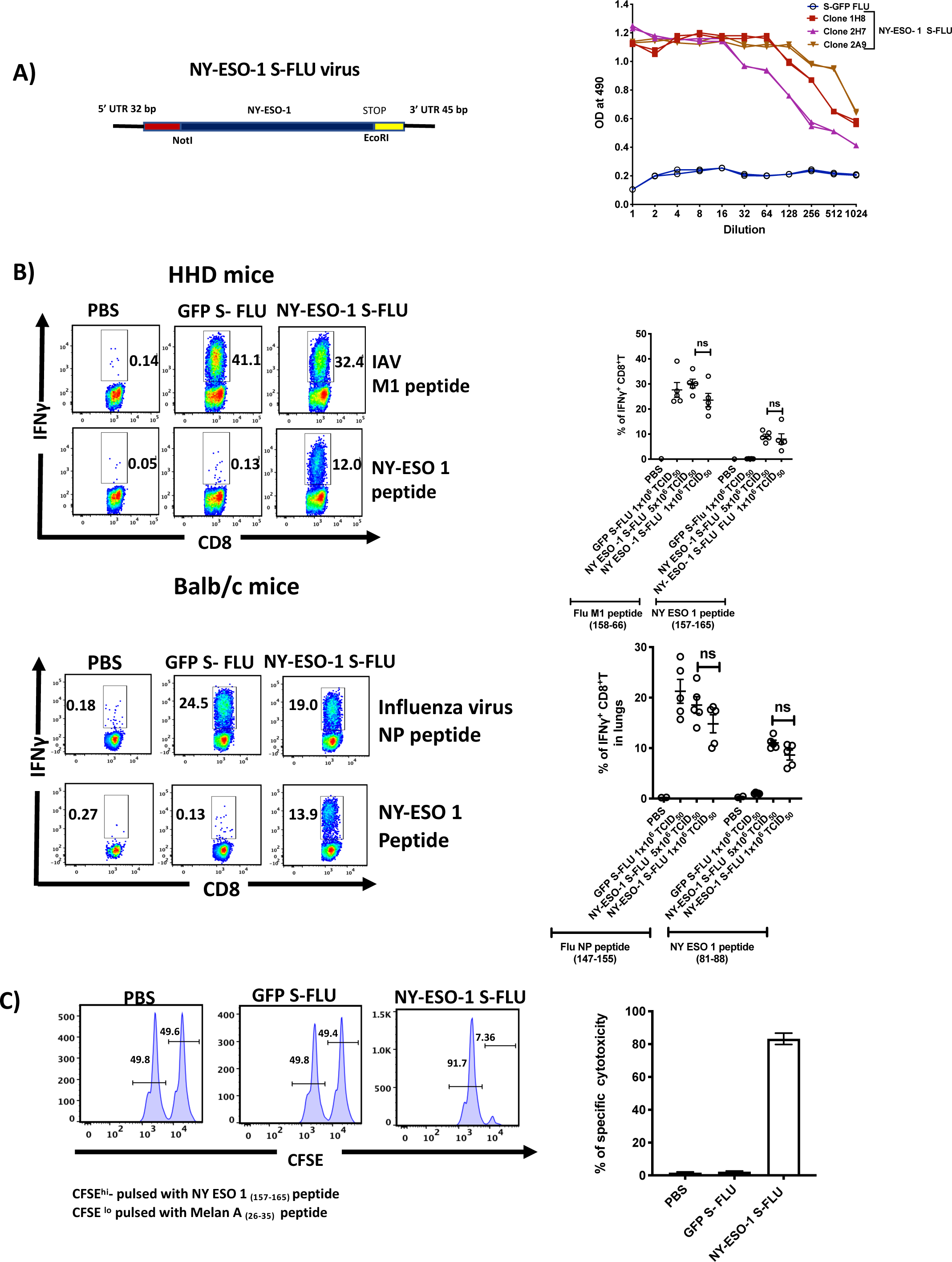
Left panel-Schematic figure of the design for NY-ESO-1 S-FLU virus. The codon optimised NY-ESO-1 cDNA sequence between unique NotI site and EcoRI sites in the pPol/S-UL expression cassette surrounded HA packaging sequences. Right panel-NY-ESO-1 expression in NY-ESO-1 S-FLU infected MDCK-SIAT1 cells. MDCK-SIA1 cells expressing HA from PR8 were infected with different clones of NY-ESO-1 S-FLU virus or GFP S-FLU virus in 2 fold serial dilutions and the expression of NY-ESO-1 was analyzed after 48hrs. Figure 1B. Left panel-Representative flow cytometry dot plots showing the frequency of IFNγ secreting CD8^+^T cells. HHD (upper panel) and Balb/c (lower panel) were intranasally infected with 1×10^6^ TCID_50_ of GFP S-FLU or 3×10^6^ TCID_50_ and 1×10^6^ TCID_50_ of NY-ESO-1 S-FLU virus on day 0. CTL responses in lungs were analyzed on day 10 post infection by ex *vivo* stimulation with HLA-A2 restricted IAV M1 peptide_158-66_(GILGFVFTL) or NY-ESO-1 peptide _157-65_(SLLMWITQC) and H-2K^d^ binding IAV NP peptide _147-55_(TYQRTRALV) or H-2D^d^ restricted NY-ESO-1 peptide _81-88_ (RGPESRLL). Right panel -bar chart shows the percentage of IFNγ secreting CD8^+^T cells in HHD (upper panel) and BALB/c mice (lower panel) lungs. Figure 1C. *In vivo* analysis of cytotoxic T cell functions. Left panel-representative FACS plots for *in vivo* killing assay. HHD mice were intranasally infected with 1×10^6^ TCID_50_ of GFP-S-FLU or NY-ESO-1 S-FLU virus and on day 10 p.i, mice were adoptively transferred with CFSE labelled (CFSE^hi^) NY-ESO-1_157-65_ peptide pulsed splenocytes and CFSE labelled (CFSE^lo^) Melan A26-35 peptide pulsed splenocytes and CFSE labelled splenocytes were analyzed in spleen after 8hrs. Right panel-Bar chart shows the percentage of NY-ESO-1 specific cytotoxicity in infected mice. The values are expressed as mean ± SEM. Data in Figure B is representative of at least two independent experiments. ns- not significant.

Next, we characterized the S-NY-ESO-1 FLU virus infection *in vitro* and *in vivo*. HEK293T cells were infected *in vitro* with S-NY-ESO-1 FLU virus in multiplicity-of-infection (MOI) 3 and the expression of NP and NY-ESO-1 were examined by immunofluorescence. Results shows that the influenza virus NP and NY-ESO-1 proteins were both expressed in infected HEK 293Tcells, with a preferential localisation within the nucleus and the cytoplasm (Supplementary figure 1). For *in vivo* infection, Balb/c mice were infected with S-NY-ESO-1 FLU virus and expression of NP was analysed in different lung cell subsets on 2 and 4 days post infection (dpi). An higher level of infection was observed on day 2 compared to day 4 p.i. Flow cytometric analysis showed that the EpCAM^+^ lung epithelial cells were the main subset of cells infected, with a low frequency of immune cells (identified as CD45^+^) positive for NP(Suppl. Fig 2).

**Figure 2.**
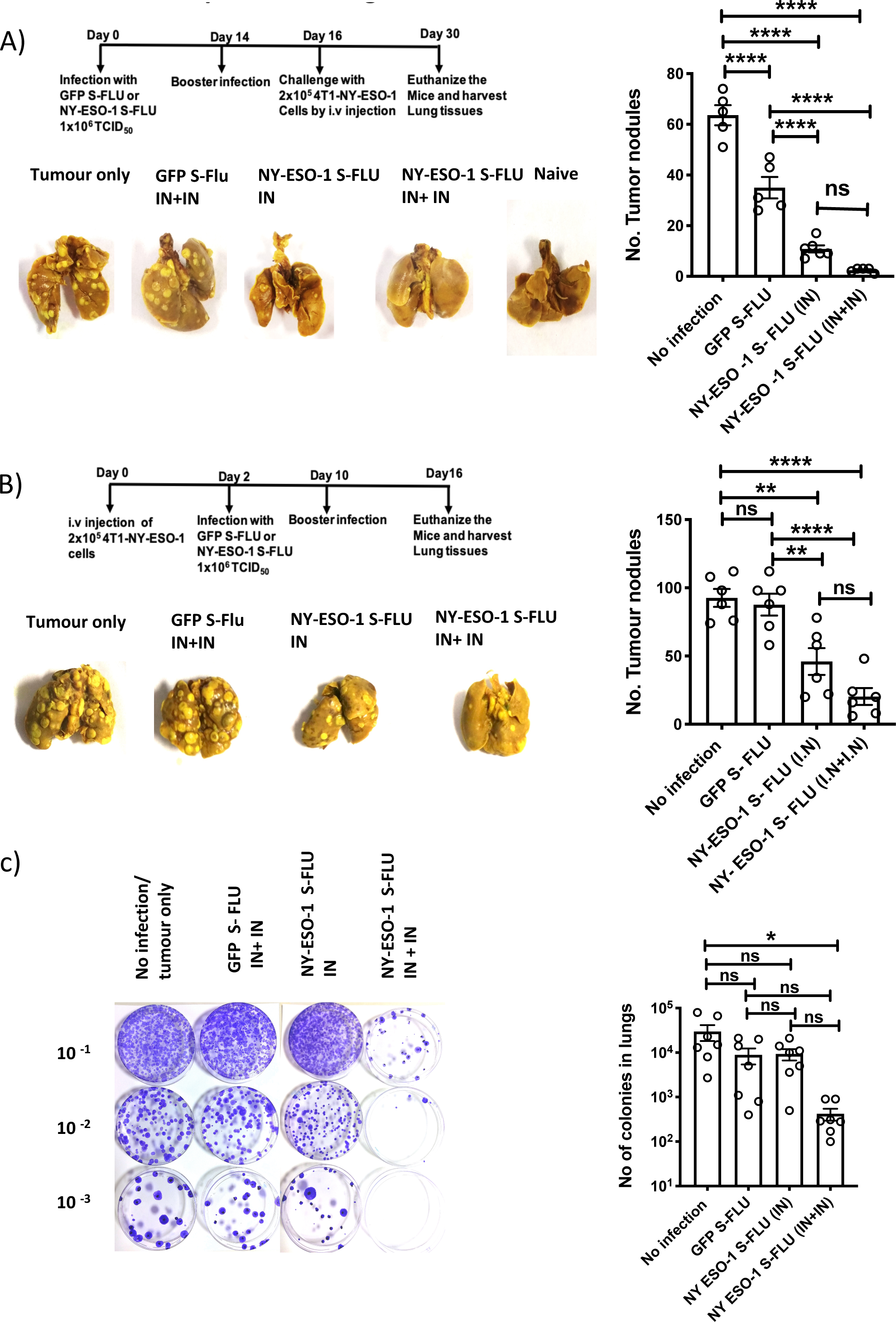
Intranasal infection with NY-ESO-1 S-FLU virus protects mice from tumour development in lungs (A) Left panel-Photographs of the lungs from the mice immunized with NY-ESO-1 S-FLU or GFP S-FLU virus followed by tumour challenge with 4-T1-NY-ESO-1 cells and right panel shows a bar chart represents the number of tumour nodules enumerated in the lungs. (B) left panel-Photographs of the lungs from the mice intravenously injected with 4-T1-NY-ESO-1 cells followed by the treatment with NY-ESO-1 S-FLU or GFP S-FLU virus and right panel shows a bar graph represents the number of tumour nodules enumerated in the lungs. Photographs are representative of 5 mice in two different experiments with similar results. (C) Clonogenic assay-Left panel - Photographs showing the spontaneous metastasis of 4T1-NY-ESO-1 cells to lungs. 4T1-NY-ESO-1 cells were injected subcutaneously in Balb/c mice and mice were intranasally infected with GFP S-FLU or NY-ESO-1 S-FLU virus on day 4 and day18 post tumour cells injection. On day 26-28 days post tumour cells injection, mice were euthanized and lung cells were plated in 6-thioguonine media. Number of colonies were counted and the right panel shows the bar chart represents the number of colonies in lungs from the mice treated with GFP S-FLU or NY-ESO-1 S-FLU virus. The results shown in (A) and (B) are a representative of two independent experiments with similar results (n = 5-6 mice/group). Data shown in (C) are pooled results of two independent experiments (n=7 mice/ group) * P<0.05, ** P<0.01, ***P<0.001 and ****P<0.0001, ns- not significant (one-way ANOVA-multiple comparisons). Error bars: mean ± SEM.

The ability of intranasal infection with S-NY-ESO-1 FLU virus to induce a T cell primary activation was then investigated in draining lymph nodes (dLN). Cell trace violet (CTV) labelled NY-ESO-1 specific CD8^+^ T cells (1G4) were adoptively transferred to HHD mice followed by infection either with GFP-S-FLU (S-FLU expressing GFP reporter protein) or NY-ESO-1 S-FLU virus and the proliferation was anlysed on day 3 p.i. Suppl Fig 3 clearly shows an effective T cell proliferation of 1G4 cells only in the groups of mice infected with the NY-ESO-1 S-FLU, and not in the control group. Next, we sought to determine if our S-FLU virus was able to induce a detectable immune response stimulating the T cell repertoire of a normal immunocompetent mouse (i.e. without any adoptive transfer). To this aim NY-ESO-1 specific CTL response was analysed in HHD (HLA-A2/Kb hybrid haplotype) or BALB/c (H-2d haplotype) mice on day 10 p.i by *ex vivo* stimulation with the relevant NY-ESO-1 CTL peptides. Fig 1B shows that intranasal infection with NY-ESO-1 S-FLU virus elicits a robust NY-ESO-1 specific CTL response in HHD (9.80±1.5%) and BALB/c mice (11.04±1.0%) albeit to a lesser magnitude than the CTL responses against the immunodominant FLU epitopes M1 and NP (30.14±3.5% and 14.82±3.6%, respectively). Intranasal infection with NY-ESO-1 S-FLU virus also induces NY-ESO-1 specific CTL response in spleen (Supplementary figure 4). The effector function of the NY-ESO-1 specific T cells was further investigated with a *in vivo* killing assay. Results showed a strong NY-ESO-1 (157-165) specific cytotoxic effect in spleens of NY-ESO-1 S-FLU infected mice compared to the control.

**Figure 3.**
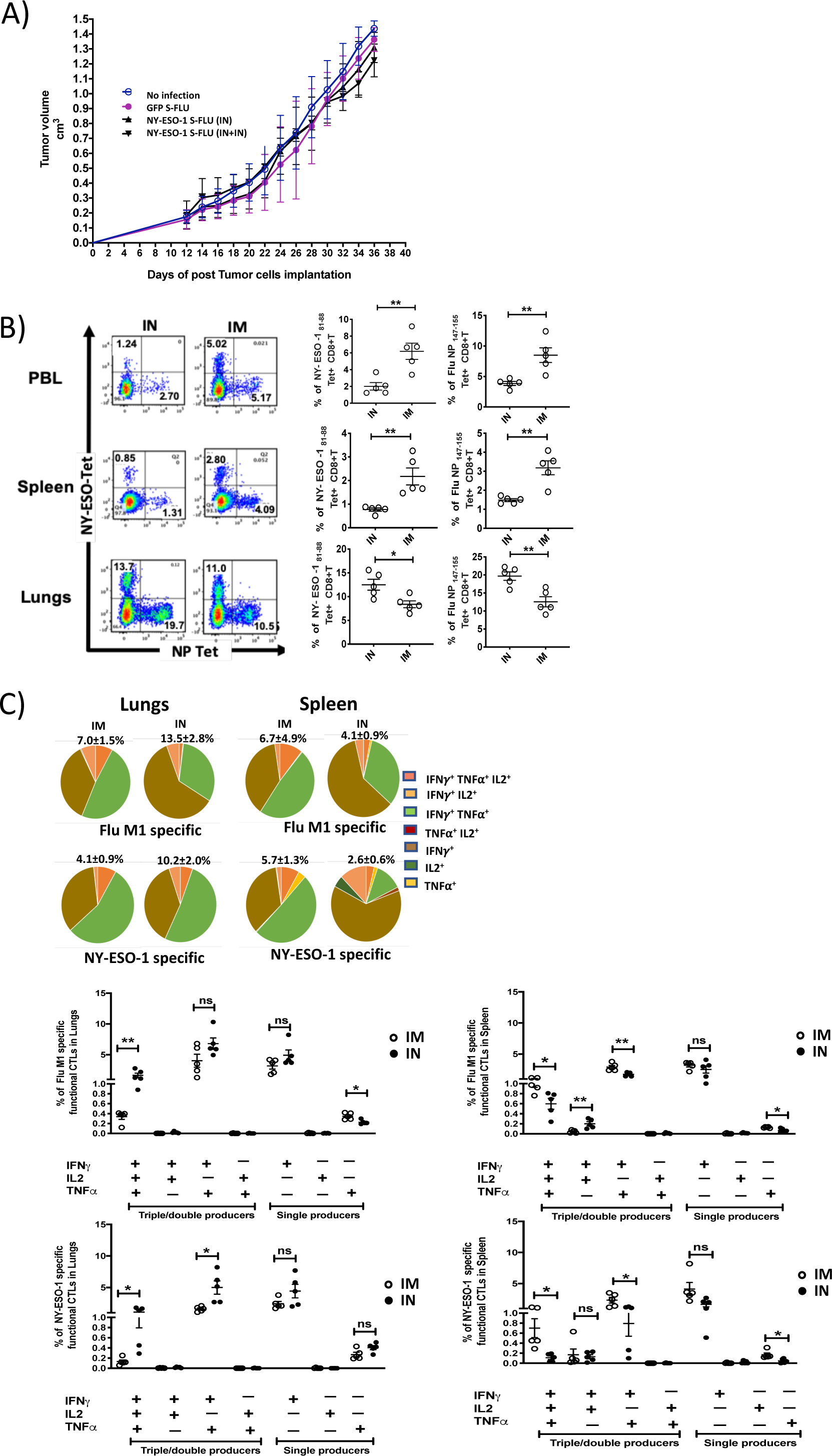
Intranasal infection with NY-ESO-1 S-FLU virus fails to protect the mice from subcutaneous tumor challenge and elicits only a modest systemic T cell response (A) Tumour growth of 4T1-NY-ESO-1 bearing mice. Balb/c mice were subcutaneously injected at right flank with 2×10^5^ cultured 4T1-NY-ESO-1 cells on day 0. On day 4 and 18, mice were intranasally infected with GFP S-FLU or NY-ESO-1 S-FLU virus. Tumour volumes were monitored every other day. (B) NP or NY-ESO-1 specific CTL responses in blood, lungs and spleen following intranasal or intramuscular infection. Balb/c mice were intranasally or intramuscularly infected with NY-ESO-1 S-FLU virus and PR8-NP or NY-ESO-1 specific CTL responses were analyzed by tetramer staining. Left panel - representative FACS plots showing the percentage of NP or NY-ESO-1 specific CD8^+^T cells in peripheral blood leukocytes (PBL), spleen and lungs. Right panel shows the bar graphs display the frequency of NP or NY-ESO-1 specific CD8^+^T cells. (C) Top panel shows the pie charts represent the frequencies of Flu M1 specific or NY-ESO-1 specific poly functional CD8^+^T cells, (triple cytokines or double cytokine producers) in lungs or spleen. HHD mice were intranasally or intramuscularly infected with NY-ESO-1 S-FLU virus and Flu M1 specific or NY-ESO-1 specific poly functional CD8^+^T cells in lungs (left) and spleen (right) were analyzed by *ex vivo* peptide stimulation on day 10 post infection. Middle panel ( Flu-M1 specific) and lower panel( NY-ESO-1 specific) show the bar graphs display the frequencies of poly functional CD8+T cells in lungs ( left) and spleen (right). The results shown in figure 3 are a representative of two independent experiments with similar results (n = 4-5 mice/group). *P<0.05, **P<0.01, ns- not significant (2 -tailed student-t test). Error bars: mean ± SEM

**Figure 4.**
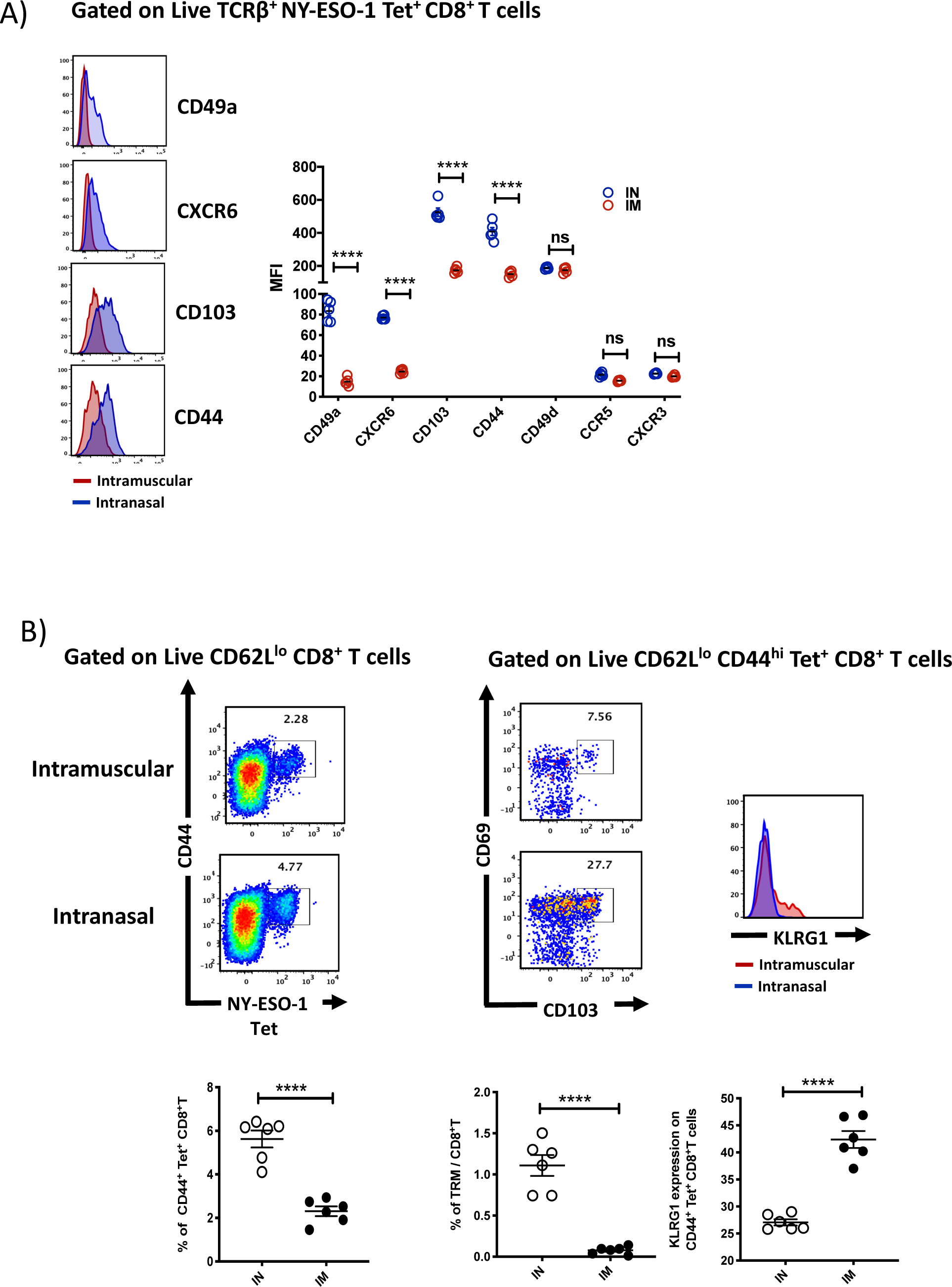
Intranasal infection induces tissue retention signals and generates a higher frequency of tissue resident memory CD8^+^T cells (A) Left panel-Representative histograms show the expression of tissue retention molecules CD49a, CXCR6, CD103 and CD44 on NY-ESO-1 tetramer positive lung CD8^+^T cells. Right panel – Bar graphs show the quantification of Mean Fluorescence Index of CD49a, CXCR6, CD103, CD44, CD49d, CCR5 and CXCR3 expressed by NY-ESO-1 specific CD8^+^T cells in lungs. Balb/c mice were intranasally or intramuscularly infected with NY-ESO-1 S-FLU virus and day 10 post infection NY-ESO-1 specific CD8^+^T cells were analyzed for various tissue retention molecules expression. The results shown are a representative of two independent experiments with similar results (n = 5-6 mice/group). (B) Compared to intranasal infection, intramuscular infection results in decreased number of tissue resident memory T cells in the lung parenchyma. Balb/c mice were intranasally or intramuscularly infected with NY-ESO-1 S-FLU virus and on day 35 p.i, the frequency of lung-resident memory CD8^+^T cells were analyzed by flow cytometry. Left panel-representative FACS plots (gated on CD8β^-^ CD8α^+^ CD62L^lo^) show CD44^hi^ NY-ESO-1 tetramer+ CD8+T cells and the FACS plots at right panel display the further analysis of CD44^hi^ NY-ESO-1 tetramer+ CD8+T cells for the expression of resident memory T cells markers-CD69 CD103 and KLRG1 (Far right histogram). Lower panel displays the bar graphs showing the percentage of tetramer positive CD8^+^T cells, tissue resident memory CD8^+^T cells among the total number of CD8^+^T cells and the percentage of KLRG1 expression on tetramer positive CD8^+^T cells. ****P<0.0001, ns- not significant (2 -tailed student-t test). Error bars: mean ± SEM

### Tumour protection following Intranasal infection with NY-ESO-1 S-FLU virus

To investigate the effectiveness of NY-ESO-1 S-FLU virus against tumour challenge, we used syngeneic 4T1 metastatic breast carcinoma (MBC) tumour model. 4T1 cells generate primary tumours and spontaneously metastasize to dLNs, lungs, spleen, liver etc. following syngeneic transplantation in immunocompetent BALB/c mice and simulates stage IV of human breast cancer progression.

We tested NY-ESO-1 S-FLU virus antitumor effect in the following settings: (1) prophylactic (2) therapeutic and (3) spontaneous tumour metastasis models. In the prophylactic model, mice were first infected with virus and subsequently challenged with 4T1-NY-ESO-1 tumour cells delivered via intravenous route. Intravenous injection of 4T1-NY-ESO-1 cells resulted in the development of tumour nodules in lungs. Mice were immunised with 1 or 2 doses of NY-ESO-1 S-FLU or with control GFP S-FLU intransally or remained naïve. Immunization with NY-ESO-1 S-FLU virus significantly reduced the number of nodules in lungs and interestingly, a single infection with NY-ESO-1 S-FLU was sufficient to give significant protection against tumour challenge (Fig 2A). For the therapeutic model, mice were intravenously injected with 4T1-NY-ESO-1 cells followed by intranasal infection with NY-ESO-1 S-FLU or control GFP S-FLU and generally a higher number of tumour nodules developed in therapeutic setting than prophylactic model (∼30 to 40%). Nonetheless, two sequential infections with NY-ESO-1 S-FLU virus significantly reduced the number of tumour nodules in lungs (Fig 2B). In the spontaneous tumour metastasis model, mice were subcutaneously injected with 4T1-NY-ESO-1 cells and subsequently infected intranasally with 1 or 2 doses of NY-ESO-1 S-FLU or GFP S-FLU or remained naive. On day 27, mice were euthanized and lungs were digested with collagenase, and dissociated cells were subjected to clonogenic assays in culture (Fig 2C). Primary tumour displayed spontaneous metastasis to lungs and mice that received two sequential infections with NY-ESO-1 S-FLU virus showed fewer colonies (421±325) (p<0.05) than any of the other treatments: no infection (29814±30716) two sequential infection with control GFP S-FLU virus (8900 ± 9215) or single infection with NY-ESO-1 S-FLU virus (9300±6916) (Figure 2C).

### Intramuscular administration induces a stronger systemic CTL response than Intranasal administration of NY-ESO-1 S-FLU virus

Despite inducing a strong CD8^+^ T cell response in lungs, intranasal infection did not elicit a high CTL response in spleen (Supplementary Fig 4). Furthermore, it failed to protect mice from subcutaneous tumour challenge with NY-ESO-1 expressing 4T1 cells (Fig 3A). A strong systemic immune response coordinated across tissues is required for tumour eradication^13^. Previously, intramuscular immunization (i.m) with recombinant adenoviral-vectored vaccine has shown generation of CTL responses in multiple mucosal sites^14^.

In order to analyse whether the i.m. route of infection with NY-ESO-1 S-FLU virus would generate a better systemic immune response, BALB/c mice were infected intranasally or intramuscularly with NY-ESO-1 S-FLU virus, and NP or NY-ESO-1 specific CD8^+^ T cell responses in peripheral blood, lung and spleen were examined by tetramer staining on day 10 post infection (Fig 3B). In lungs, intranasal infection induced a significantly higher NP (p<0.01) and NY-ESO-1 (p<0.05) specific CD8^+^T cell response compared with intramuscular injection. In the blood and spleen, intramuscular infection induced stronger NP (p<0.01) and NY-ESO-1 (p<0.01) specific CTL responses than intranasal infection (Figure 3B). Similarly, an increased frequency of Flu M1 or NY-ESO-1 specific IFNγ ( single producer), or IFNγ and TNFα (double producer) or IFNγ, TNFα and IL2 producing CD8^+^ T cells (triple producers) was observed in HHD mice spleen following intramuscular infection (Figure 3C) and conversely a higher frequency of flu M1 or NY-ESO-1 specific IFNγ or IFNγ and TNFα producing CD8^+^ T cells were detected in HHD mice lungs following intranasal infection (Figure 3C). Moreover, intranasal infection and intramuscular infection resulted in an accumulation of higher number of total Flu M1 / NY-ESO-1 specific CD8^+^ T cells in lungs and spleen respectively (Supplementary Fig 5).

**Figure 5.**
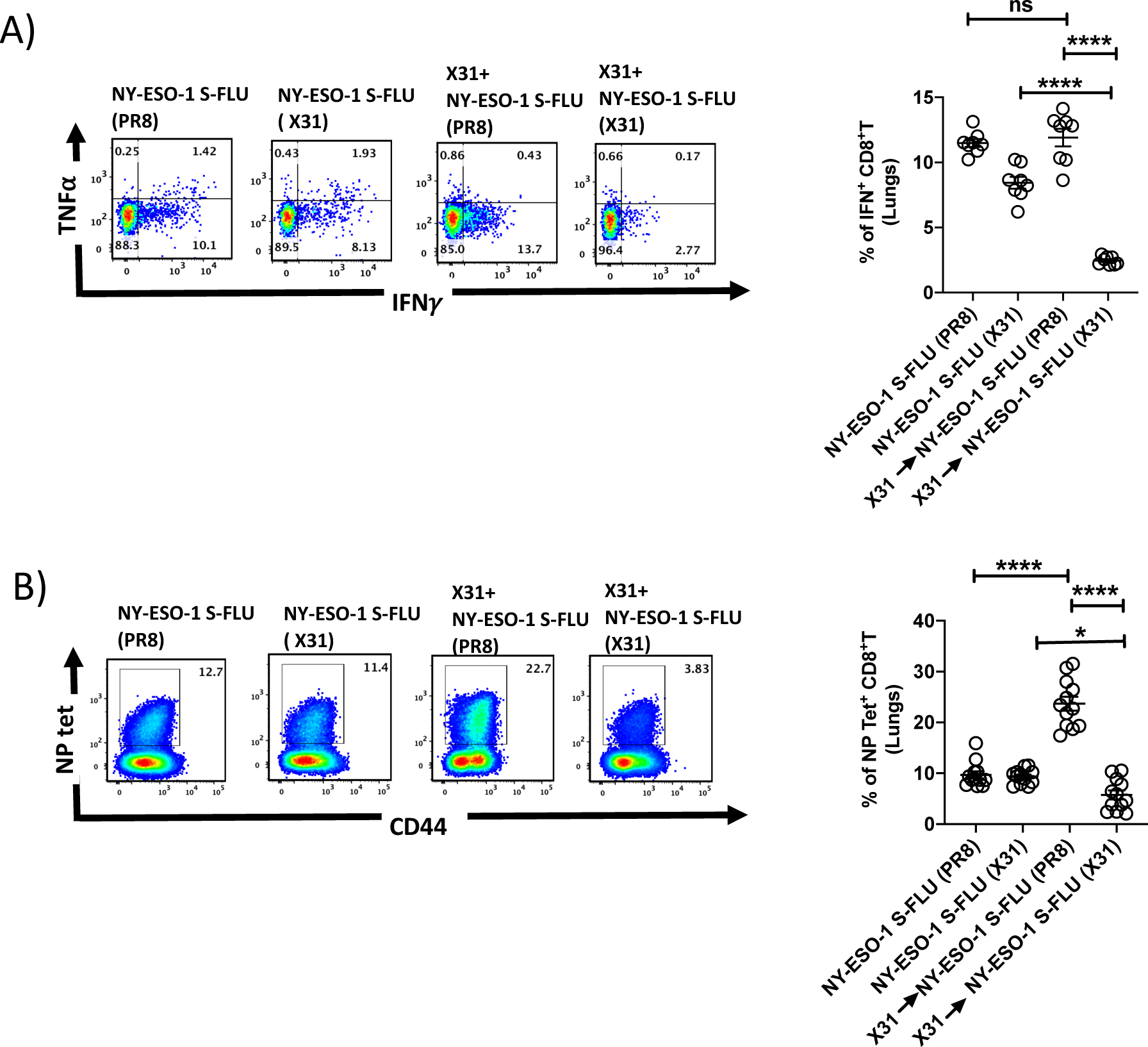
Infection with HA switched NY-ESO-1 S-FLU virus overcomes the inhibitory effect of pre-existing neutralizing antibody (A-left) Representative FACS plot showing IFNγ and TNFα secreting and (B-left) NP tetramer positive CD8^+^T cells in lungs. Bar graph show the quantification of IFNγ secreting CD8^+^T cells (A-right) and NP tetramer positive CD8^+^T cells (B-right) in lungs. BALB/c mice were intranasally infected with X31 virus and re-infected with NY-ESO-1 S-FLU (X31) or NY-ESO-1 S-FLU (PR8) on day 24 p.i and NY-ESO-1 and NP specific CD8^+^T cell responses in lungs were analyzed on day 7 post-secondary infection by *ex vivo* stimulation with NY-ESO-1 CTL peptide _81-88_ (RGPESRLL) and NP tetramer staining respectively. Data shown in (A) and (B) are pooled results of two (n=8 mice/group) and three independent experiments (n = 12 mice/group) respectively. *P<0.05, and ****P<0.0001 and ns- not significant (one-way ANOVA-multiple comparisons). Error bars: mean ± SEM.

### Intranasal infection induces the expression of tissue retention signals and demonstrates a distinct phenotype

The reciprocal distribution of antigen specific CD8^+^ T cells in lungs and spleen in post intranasal infection indicates that the effector T cells are mostly retained in lungs following intranasal infection. Activated antigen specific CD8^+^ T cells have been known to persist following recovery from respiratory virus infection and Very Late Antigen 1 (VLA-1; α1β1 integrin) has been implicated in the retention of T cells ^15–17^.

We therefore investigated whether the intranasal or intramuscular infection with NY-ESO-1 S-FLU virus differentially induced the expression of the molecules implicated in retention or trafficking of effector CD8^+^ T cells in lung. HHD mice were infected intranasally or intramuscularly with NY-ESO-1 S-FLU virus and the lung cells were analysed for the expression of VLA-1 (CD49a), CXCR6, CD103, CD44, CD49d, CCR5 and CXCR3 on day 10 post infection. The expressions of VLA-1, CXCR6, CD103 and CD44 (p<0.0001) were significantly upregulated on NY-ESO-1 specific CD8^+^ T cells in the lungs of intranasally infected mice (Figure 4A). Furthermore, the frequencies of tissue resident memory CD8^+^ T cells (TRM) (p<0.0001) were also significantly higher in intranasally infected mice (Figure 4B)

### Swapping the coat HA in NY-ESO-1 S-Flu virus overcomes the inhibitory effect of pre-existing neutralizing antibodies

Pre-existing antibodies derived from previous influenza infection or vaccination could interfere with vaccine immunogenicity and potentially impact vaccine efficiency. NY-ESO-1 S-FLU virus can be pseudotyped with any HA in the envelope and coating with relatively novel HA against whose antibodies are scarcely presented in major population could nullify the pre-existing antibody mediated effect. To validate that hypothesis, mice were first infected via the intranasal route with X31 (H3N2) virus to elicit the antibody (against H3N2) production and on day 24 p.i, mice were re-infected with NY-ESO-1 S-FLU virus with matched HA/NA (NY-ESO-1 S-FLU (X31)) or mismatched HA/NA (NY-ESO-1 S-FLU (H1 PR8)). NY-ESO-1 specific CD8^+^T cell responses in lungs or spleen were analyzed on day 7. NY-ESO-1 and NP specific CTL responses were reduced in lungs following the infection with NY-ESO-1 S-FLU virus with matched HA/NA (Figure 5A&B). Similarly in spleen, infection with NY-ESO-1 S-FLU virus with different HA coating (NY-ESO-1 S-FLU (PR8)) elicited stronger NP specific CTL response (p<0.0001). However, NY-ESO-1 specific CTL response in spleen was reduced in both matched or mismatched HA/NA NY-ESO-1 S-FLU virus infections (Supplementary Figure 7A&B)

### Intramuscular injection of NY-ESO-1 S-FLU virus induces a higher recruitment of NY-ESO-1 specific CD8^+^T cells at tumour site and reduces tumour burden

To evaluate the effectiveness of intramuscular infection in tumour bearing mice, mice with 4T1-NY-ESO-1 established subcutaneous tumour were intramuscularly or intranasally administered with NY-ESO-1 S-FLU virus. Tumour size was measured, and mice were euthanized when the tumour volume reached humane end point. NY-ESO-1 specific CD8^+^ T cells were analysed in tumour, lungs and spleen at the end time point. Mice treated with intramuscular injection showed a greater reduction in tumour size (p<0.05 on day 18,20,24 and (p<0.01 on day 26 and 28) which was concomitant with significantly higher infiltration of NY-ESO-1 specific CD8^+^ T cells in TIL (p<0.01) (Figure 6A) whereas, the mice infected via the intranasal route did not show any difference in tumour size (Figure 6A). NY-ESO-1 specific CTLs that infiltrate the tumour express a higher level of PD1. We also investigated the distribution of NY-ESO-1 specific CD8^+^ T cells in lungs and spleen and strikingly a higher number of NY-ESO-1 specific CD8^+^ T cells still accumulated in lungs in intranasally infected mice. However, there was no difference in the distribution of NY-ESO-1 specific CD8^+^ T cells splenocytes) infected mice (Figure 6B).

**Figure 6.**
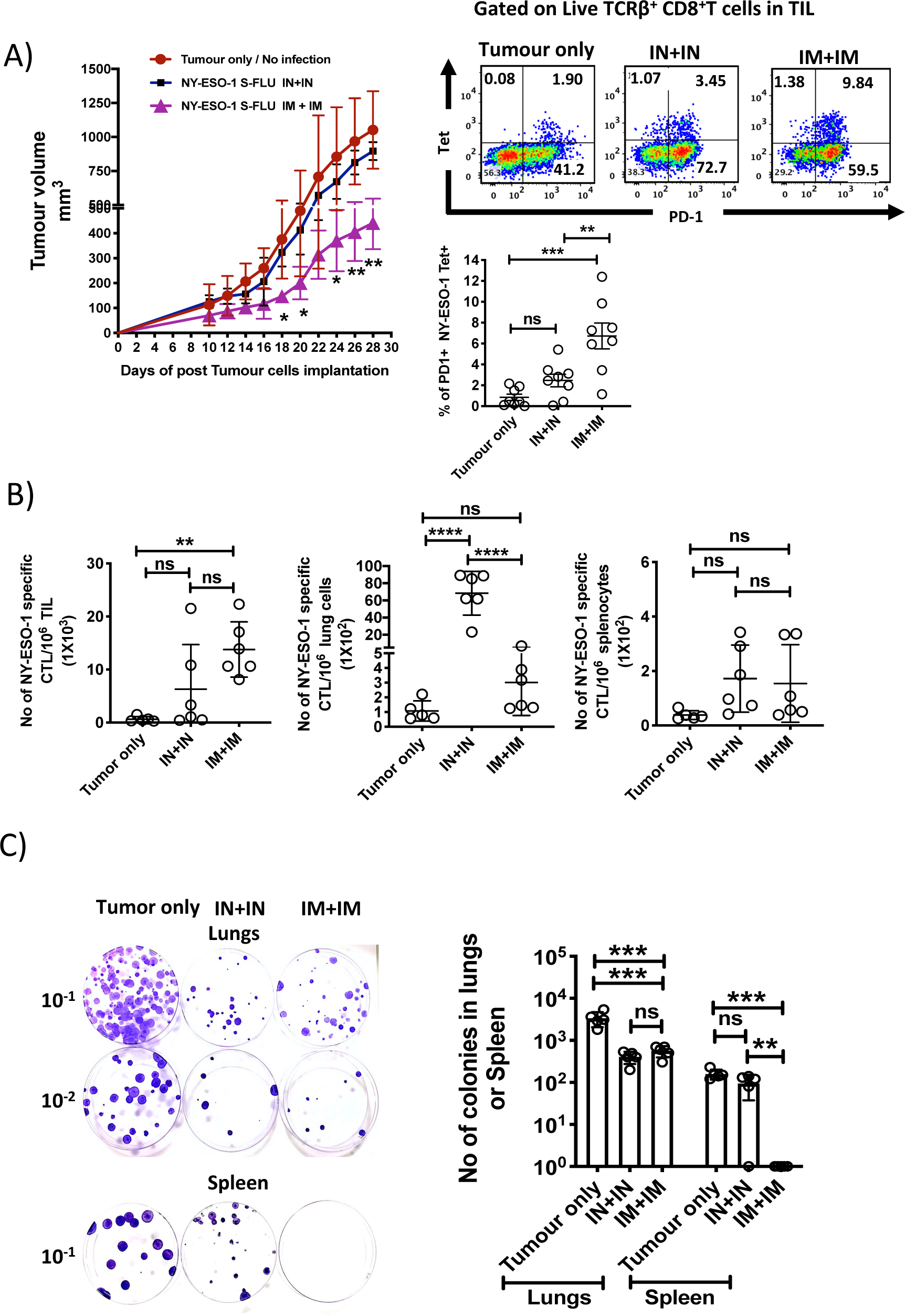
Intramuscular injection of NY-ESO-1 S-FLU virus induces a higher infiltration of NY-ESO-1 specific CD8^+^T cells at tumor site and reduces Tumor burden (A) Intramuscular infection with NY-ESO-1 S-FLU virus reduces tumour burden. Balb/c mice were subcutaneously injected with 4T1-NY-ESO-1 cells and day 4 post injection mice were intranasally or intramuscularly infected with NY-ESO-1 S-FLU virus followed by booster infection on day 18. Tumour growth was monitored over time. Tumour volume measured 28 days post inoculation in uninfected Vs intranasal infection Vs intramuscular infection is shown (left panel). Right panel shows the representative FACS plots of PD-1^+^ NY-ESO-1 tetramer^+^ CD8^+^T cells infiltrated to tumour. Bar graph shows the frequencies of PD-1^+^ Tet^+^ CD8^+^T cells in tumour infiltrating leukocytes (TIL) in the mice received no infection or intranasal or intramuscular infection with NY-ESO-1 S-FLU virus. (B) Bar graphs show the number of NY-ESO-1 tetramer+ CD8+T cells in TIL or lungs or spleen in tumour bearing mice received no infection or intranasal or intramuscular infection. (C) Clonogenic assay-Photographs (left) showing the colonies which represent the spontaneous metastasis of 4T1-NY-ESO-1 cells to lungs. On day 28 post tumour cells inoculation, mice were euthanized followed by digestion of lungs and spleen and the cells were plated in 6-thioguonine media. Number of colonies were counted after 14 days of incubation. The bar chart (right) represents the number of colonies in lungs and spleen. The results shown in figure 5 are representative of two independent experiments with similar results (n = 4-6 mice/group). Data shown in bar graph in (A) are pooled results of two independent experiments (n=8 mice/ group). * P<0.05, ** P<0.01, ***P<0.001 and ****P<0.0001, ns- not significant (one-way ANOVA-multiple comparisons). Error bars: mean ± SEM.

To determine whether the infection through different routes would induce a diverse protection against spontaneous metastasis, from a primary subcutaneous tumour to the lungs, mice with 4T1-NY-ESO-1 established subcutaneous tumour were intramuscularly or intranasally administered with NY-ESO-1 S-FLU virus. At humane end point, mice were euthanized and lung or spleen homogenates were tested in clonogenic assay. In clonogenic assay, the number of colonies in lungs following intranasal or intramuscular infection were not significantly different from each other but lower than the mice with no infection (p<0.001) (Figure 6C). However, the impaired metastases in spleen was only observed in intramuscularly infected mice as depicted in Figure 6C.

### Intramuscular injection with NY-ESO-1 S-FLU virus induces a stronger antitumor response than other contemporary virus based vaccines

Several viruses have been exploited as vehicles for delivering cancer antigens and we analysed the CTL responses induced by the widely employed or clinically tested viral vectors, based on Adenovirus ^18, 19^ and Fowl Pox virus ^9, 20, 21^ that express full length NY-ESO-1 protein. Mice were intramuscularly infected with 5×10^7^ plaque forming unit (PFU) NY-ESO-1 expressing Fowl Pox virus (Fowl Pox-NY-ESO-1) or 1×10^9^ PFU human adeno virus 5 (Hu-Ad5-NY-ESO-1) or 6×10^6^ TCID_50_ S-NY-ESO-1 FLU virus and NY-ESO-1 specific CTL responses were analysed in spleen and lungs. Infections with recombinant NY-ESO-1 S-FLU and Hu-Ad5-NY-ESO-1 viruses display comparable CTL responses in lungs and spleen whereas Fowl Pox infection did not elicit a stronger NY-ESO-1 specific CTL response (Supplementary Fig 8). To evaluate anti-tumour response, mice with established subcutaneous tumour as described above were intramuscularly injected with either NY-ESO-1 S-FLU, Hu-Ad5 NY-ESO-1, or Fowl Pox-NY-ESO-1. Mice treated with NY-ESO-1 S-FLU virus infection displayed a reduced tumour growth (716.9 ± 125.8 mm^3^) compared to the mice treated with Hu Ad5-NY-ESO-1 (972.0 ± 90.6 mm^3^) or Fowl Pox-NY-ESO-1 (1137.4 ± 223.6 mm^3^) viruses. (Figure 7A).

**Figure 7.**
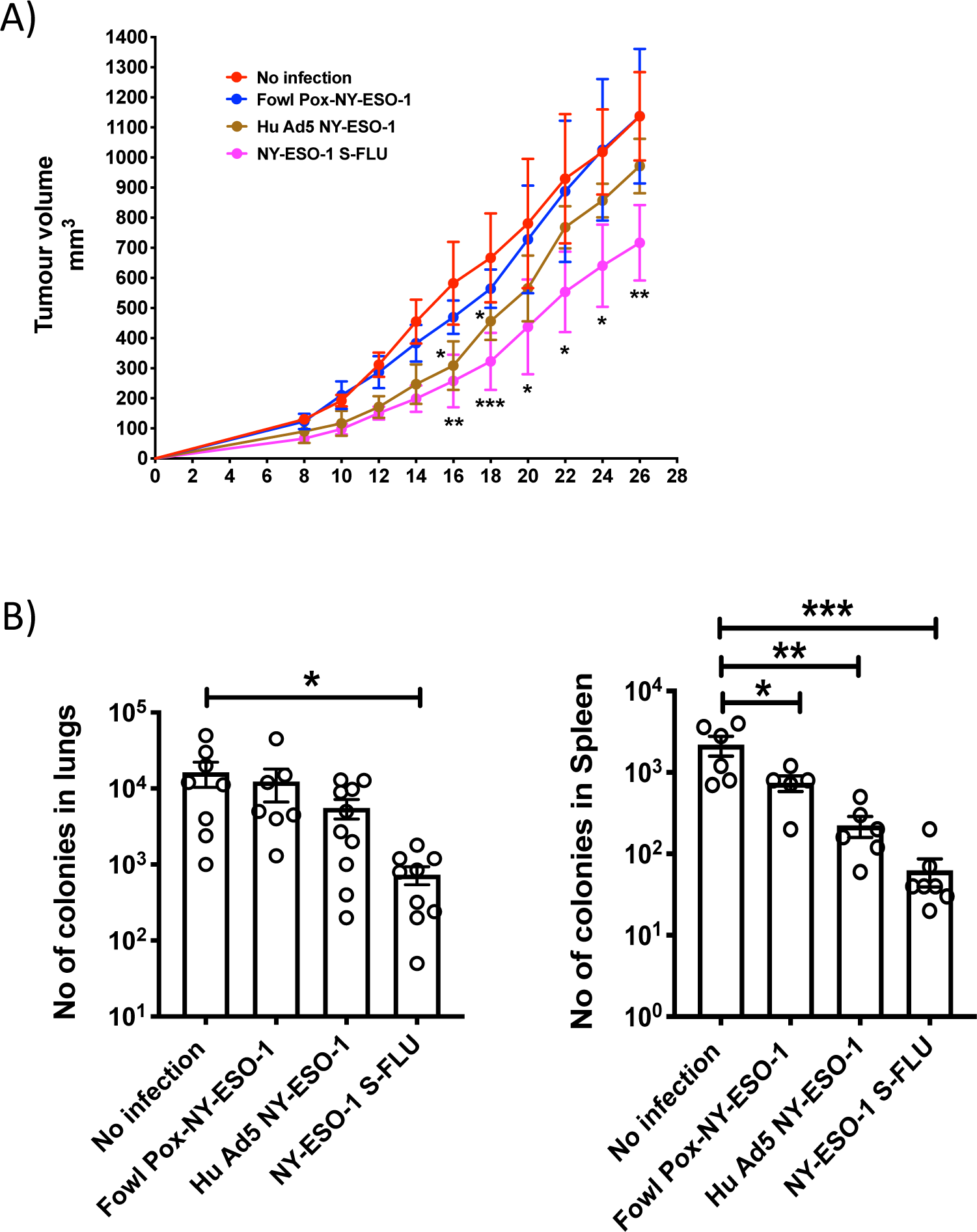
Comparison of the anti tumour effect induced by NY ESO-1 S-FLU virus with contemporary recombinant virus vaccines (A) Intramuscular injection with NY-ESO-1 S-FLU virus demonstrates a better tumour reduction than Hu Ad5 NY-ESO-1 virus or Fowl Pox NY-ESO-1 virus. Balb/c mice were subcutaneously injected with 4T1-NY-ESO-1 cells and day 4 post injection mice were intramuscularly infected with 5×10^7^ PFU NY-ESO-1 expressing Fowl Pox virus or 1×10^9^ PFU Human Adeno virus 5 or 6×10^6^ TCID_50_ NY-ESO-1 S-FLU virus followed by same dose booster infection on day 18. Tumour growth was monitored over time. Tumour volume measured 28 days post inoculation in uninfected Vs NY-ESO-1 S-FLU Vs Hu Ad5 NY-ESO-1 Vs Fowl Pox NY-ESO-1 virus infection. (B) Clonogenic assay-Bar graphs showing the number of colonies which represent spontaneous metastasis of 4T1-NY-ESO-1 cells to lungs (left) and spleen (right) in different virus infections described in (A). * P<0.05, ** P<0.01, and ***P<0.001 and ns- not significant (one-way ANOVA-multiple comparisons). Error bars: mean ± SEM.

**Figure 8.**
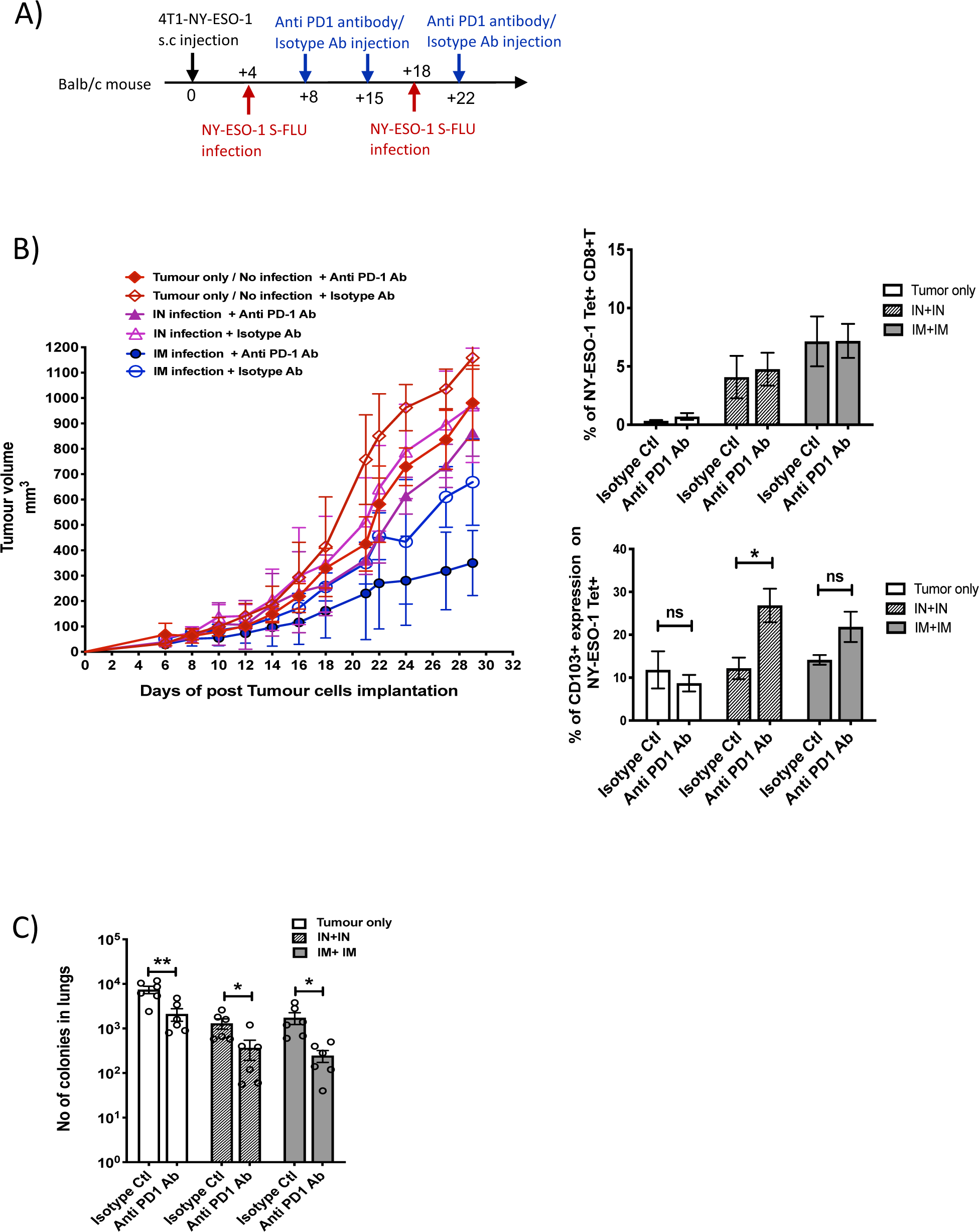
Blockade of PD-1 augments the antitumor effect of NY ESO-1 S-FLU virus infection (A) Experimental design. (B) Tumour growth curves from the experiment described in (B) (left) and bar graphs show the frequency of NY-ESO-1 tetramer^+^ CD8^+^T cells in TIL (right upper) and the percentage of CD103 expression on NY-ESO-1 tetramer+ CD8+T cells in TIL (right lower). (C) Clonogenic assay-Bar chart shows the number of colonies in lungs. The results shown in figure 8 are representative of two independent experiments with similar results (n = 5-6 mice/group). *P<0.05, **P<0.01, ns- not significant (2 -tailed student- t test). Error bars: mean ± SEM

Moreover, mice that received NY-ESO-1 S-FLU virus demonstrated a lower number of 4T1-NY-ESO-1 clones derived from metastatic niches at lungs or spleen in clonogenic assay compared to the mice treated with Hu Ad5 NY-ESO-1 or Fowl Pox virus infection (Figure 7B).

### Blockade of PD1 augments the antitumor effect of NY-ESO-1 S-FLU infection

We observed that cytotoxic T cells infiltrated to the tumour displayed a higher expression level of PD1 (Figure 6A). To test the possible immune checkpoint role of PD-1 in this setting, and to improve the effectiveness of NY-ESO-1 S-FLU virus mediated anti-tumour responses, a combined treatment of NY-ESO-1 S-FLU virus and anti PD1 antibody was tested. Anti PD1 was chosen as it is a widely used for check point blockade (CPB) and also an approved drug for many cancer conditions. In general, 4T1-NY-ESO-1 tumours positively responded to both anti PD1 monotherapy and combined therapy with NY-ESO 1 S-FLU virus infection by displaying a slower tumour progression compared to isotype antibody treatment (Figure 8B left panel). The most pronounced tumour regression was observed in combined treatment with anti PD1 antibody and intramuscular injection with NY-ESO-1 S-FLU virus (Figure 8B left panel). The anti PD1 antibody administration did not increase infiltration of the NY-ESO-1 specific CD8^+^T cells into the tumour in NY-ESO-1 S-FLU virus infected mice (Figure 8B right panel), but it was associated with increased expression of CD103 on NY-ESO-1 specific CTLs in TIL (Figure 8B right panel).

Next, we investigated whether the combined treatment of anti PD1 antibody and NY-ESO-1 S-FLU virus infection could display an augmented inhibition of spontaneous metastasis to lungs. As depicted in Figure 8C, anti PD1 antibody monotherapy showed a reduced tumour metastasis to lungs and notably, in combined therapy with NY-ESO-1 S-FLU virus infection it demonstrated a significantly higher reduction in tumour metastasis (p<0.05).

## Discussion

The major challenge for the development of tumour vaccines is identifying the delivery system that is capable of inducing safer and more efficient T cell responses. Recombinant influenza viruses expressing TAA were shown to reduce the tumour burden through the induction of cytotoxic T-cell responses and therefore makes it an attractive tumour antigen delivery vector ^22, 23^. Among many types of influenza vaccines, live attenuated influenza vaccine (LAIV) is one of the strongest inducers of CD8^+^T cell response ^24^ and the S-FLU virus used in this study is similar to LAIV in terms of generating T cell mucosal immunity in mice and ferrets^11, 25^. We investigated the vaccine efficacy of nonreplicating S-FLU vaccine in inducing specific CTL responses and inhibiting tumour development and progression. Intranasal immunization of NY-ESO-1 S-FLU virus induced a robust specific CTL response in lungs but with a modest systemic CTL response in spleen. The route of vaccine administration influences the intensity of the systemic antigen specific T cell response. H5N1 whole inactivated virus (WIV) immunization via intranasal and intramuscular route induced a comparable frequency of multifunctional Th1 CD4^+^ cells (IFN-γ, IL-2 and TNF-α) in spleen ^26^ whereas PR8 WIV strain infection induced a higher IFNγ secreting CD4^+^ T cells in spleen^27^ only in intranasal route. We observed a significantly higher NY-ESO-1 specific CD8^+^T cells in blood and spleen with intramuscular immunization which is in line with a previous study^28^.

An orthotopic 4T1 mouse breast cancer model used in this study resembles triple-negative breast tumours (ER^-^, PR^-^, HER2^-^) in humans^29^. Intranasal infection with NY-ESO-1 S-FLU virus protected the mice from lung metastasis induced by tail vein or subcutaneous injection. Interestingly, S-FLU without NY-ESO-1 showed a partial but significant protection against tumour development in lungs (Figure 2A), suggesting that nonspecific innate immune response mediated through TLR7 by virus single strand RNA may be utilized for anti-tumour response^30^. Nonetheless, full reduction of spontaneous metastasis was only achieved with NY-ESO-1 S-FLU virus infection, but without any impact on subcutaneous tumour development. Regional tumour protection with intranasal infection may be attributed to the continued presence of antigen specific CTLs in lungs which was associated with poor infiltration of CTLs into tumour. Recently, it has been shown that intranasal infection with influenza virus accelerated melanoma growth in skin with increased shunting of anti tumour CD8^+^T cells from the tumour site (skin) to distant site (lungs) resulting in decreased immunity within the tumour ^31^. Moreover, increased upregulation of tissue retention molecules VLA-1, CXCR6, CD103 and CD44 on lung CD8^+^T cells following intranasal infection (Figure 4(A)), probably prevents the T cells migration to distal subcutaneous tumour site, thus permitting the uninhibited tumour growth.

In contrast, intramuscular injection with NY-ESO-1 S-FLU virus reduced primary tumour burden with decreased spontaneous tumour metastasis to lungs and spleen (Figure 6C). Intriguingly, primary tumour was highly infiltrated with antigen specific CD8^+^T cells but not in lungs or spleens which suggests that decreased metastasis was likely due to primary tumour regression.

As reported before ^32^ CTL responses induced with S-FLU virus and recombinant adenovirus infection were comparable (in lungs and spleen); however, further investigations is needed to understand whether both viral infections induce a similar frequency and quality (poly functionality or tumour cell killing effect) of tumour infiltrating antigen specific CD8+T cells following infection. With a booster immunization, S-FLU virus induced a more efficient tumour regression than recombinant adenovirus (Figure 7). The weaker efficacy in tumour reduction with adenovirus may be due to anti-vector neutralizing antibodies raised from primary infection^33^ which hampers the booster immunization effect. It has also been highlighted recently that the cellular immune response was not further enhanced following booster immunization with chimpanzee adenovirus-vectored vaccine (ChAdOx1 nCoV-19) for SARS-CoV-2 ^34^ and options like using different adenoviral vectors for the booster immunization or extending the time more than 10 months between two inoculations have been suggested to overcome fallibility^35^. Pre-existing influenza virus immunity may also limit the replication and immunogenicity of influenza based viral vaccine. However, this can be overcome by pseudotyping S-NYE-SO-1 FLU with a novel HA as shown in the proof of principle experiment in Figure 5.

Despite a significant tumour reduction over intramuscular infection, tumour progression was not completely impaired suggesting an immune suppressive tumour micro environment which is substantiated with the higher expression levels of PD-1 on cytotoxic T cells in tumour (Figure 6A). Suppression of T cell function is mediated through PD-1 signalling with its ligands PD-L1 and PD-L2 in murine ^36^, patient tumours ^37^ and the successful reduction of the tumours has been achieved with check point inhibitors ^29 38–41 42^. In our study, treatment with anti PD-1 antibody showed a modest tumour reduction on its own, while the administration in combination with intramuscular delivery of NY-ESO-1 S-FLU virus displayed a synergistic effect with a drastic reduction in tumour size. We administered PD-1 antibody three times (Figure 8A) since >4 times injection was known to cause fatal hypersensitive reaction in 4T1 tumour bearing mice^43^ and local delivery of PD-1 antibody may reduce systemic adverse effect and would allow more frequent administration. In a combination therapy of intramuscular infection with a subsequent PD-1 antibody administration, CD103 expression was upregulated on cytotoxic T cells and it is correlated with tumour reduction as CD103 on CTLs improves TCR antigen sensitivity, enables faster cancer recognition and rapid antitumor cytotoxicity ^44, 45^. It would be interesting to investigate in future whether CD103 expression on CTLs is associated with E-cadherin or ICAM-1 expression on tumour cells and strong adhesion between the molecules expressed by tumour cells and CD103 on CTLs requires for efficient tumour reduction^46^.

In conclusion, this study outlines the potential of S-FLU virus as a tumour vaccine that drives the generation of stronger antigen specific CTL response and dissimilar immune responses elicited in intranasal and intramuscular administration. NY-ESO-1 S-FLU virus induced antitumour responses occurred regionally (lungs-intranasal administration) and systemically (spleen and blood-intramuscular administration), were tumour antigen specific, and were associated with T cell infiltration in tumours. Moreover, in combination with PD-1 blockade NY-ESO-1 S-FLU delivery results in drastic reduction in tumour growth. Our study suggests that an S-FLU based tumour vaccine shows great promise for tumour protection in lungs via intranasal administration and for better protection against peripheral tumour via intramuscular administration. In future studies, we would like to investigate whether the synergy of vaccinations with NY-ESO-1 S-FLU virus (intramuscular) and PD-1antibody expressing S-FLU (intratumoural injection) could clear the tumour completely ^47 31^.

## Methods

### Mice

Human leukocyte antigen (HLA)-A2.1 transgenic mice ^48^ (HHD mice), and 1G4 transgenic mice expressing the HLA-A2/NY-ESO-1_157-165_ specific TCR (^49^ were bred in the local animal facility under specific pathogen–free conditions and used at 6 to 10 weeks of age. 6-7 weeks old BALB/c mice were purchased from Envigo. Animal studies have been conducted in accordance with, and with the approval of, the United Kingdom Home Office. All procedures were done under the authority of the appropriate personal and project licenses issued by the United Kingdom Home Office License number PBA43A2E4.

### Cell culture and recombinant viruses generation

HEK, 4T1 and MDCK-SIAT1 cells used in this study were obtained from ATCC. HEK and MDCK-SIAT1 cells were maintained in DMEM, 4T1 cells were maintained in RPMI (Gibco). Media were supplemented with 10% fetal bovine serum (FBS), 2 mM Glutamine and penicillin/streptomycin. NY-ESO-1 S-FLU was generated as previously described (Powell et. al., 2012) with minor modification. In brief, the codon optimised cDNA encoding NY-ESO-1 flanked by *NotI* and *EcoRI* cloning sites was synthesised by GeneArt and ligated into the modified S-FLU expression plasmid pPol/S-UL between the 3’ and 5’ HA packaging sequences. The inactivated HA signal sequence (part of the packaging signal) was optimised by removal of unwanted ATG sequences. GFP S-FLU was made similarly with GFP replacing the NY-ESO-1 sequence. Recombinant NY-ESO-1 S-FLU on the A/PR/8/1934 background were produced by transfection of HEK 293T cells as described ^11, 50^ and cloned twice by limiting dilution in MDCK-SIAT1 cells stably transfected to express coating haemagglutinin from A/PR/8/34 [GenBank accession no. CAA24272.1] to provide the pseudotyping haemagglutinin in *trans* in viral growth media (VGM; DMEM with 1% bovine serum albumin (Sigma-Aldrich A0336), 10 mM HEPES buffer, penicillin (100 U/ml) and streptomycin (100 ug/ml)) containing 0.75ug to 1ug/mL of TPCK-Trypsin (Thermo Scientific, 20233). The NY-ESO-1 S-FLU (PR8) was harvested after 48 hours by centrifugation (1,400g for 5 min) of the culture supernatant to remove debris and kept as a seed virus. NY-ESO-1 S-FLU (X31) was generated by infecting MDCK-SIAT1 stably transfected with X31 H3 (MDCK-X31) with the NY-ESO-1 S-FLU (PR8) seed virus (∼MOI 0.01) and the supernatant was harvested as described above after 48 hours. NY-ESO-1 S-FLU was titrated as TCID_50_ as previously described (Powell et al., 2012, Powell et al., 2019). In brief, supernatant containing NY-ESO-1 S-FLU were titrated in 1/2-log serial dilution in VGM (total 50uL) across a flat-bottom 96-well plate seeded with 3e4 MDCK-PR8 or MDCK-X31 cells. After 1 hour incubation at 37C, 150uL of VGM containing 1ug/mL TPCK-Trypsin was added into each well and the plate was incubated at 37C for 48 hours. After, that, plates were washed twice with PBS then fixed with 100 ul of 10% formalin in PBS for 30 min at 4C and permeabilised with 50 ul of permeabilization buffer (PBS, 20mM glycine, 0.5% Triton-X100) for 20min at RT. Plates were then washed twice with PBS and stained with 50 uL of PBS containing 0.1% BSA (PBS/0.1% BSA) and mouse anti-NY-ESO1 antibody (Cat no, 1:250) for 1 hour. Plates were then washed twice with PBS and stained with 50 uL goat-anti-human AF647 for 1 hour. Plates were then washed with PBS twice and fluorescence was measured using a ClarioStar Plate Reader (BMG Labtech). TCID_50_ was calculated using the Reed and Muench method (Reed and Muench, 1938).

The full-length NY-ESO-1 gene (Genbank No: NM001327) was cloned into the replication deficient human adenovirus serotype 5 (AdHu5) vector backbone under the control of the human CMV immediate early long promoter to generate the HuAd5-NY-ESO-1 construct. The replication-deficient adenoviral vectors were scaled up by the Viral Vector Core Facility at the Jenner Institute (Oxford, UK) in 293A cells with purification by caesium chloride centrifugation and stocks were stored at -80**°**C in PBS. Virus titre was determined in a cytopathic effect assay^51^. Purity and sterility were confirmed by PCR and inoculation in TSB broth, respectively.

### Virus infections

Influenza virus infections via intranasal route (50 μl volume) were performed with 1×10^6^ TCID_50_ of GFP S-FLU, 1×10^6^ or 3×10^6^ TCID_50_ of NY-ESO-1 S-FLU (PR8) virus or NY-ESO-1 S-FLU (X31) virus and 3.2×10^4^ TCID_50_ of X31 virus. For intramuscular route, 6×10^6^ TCID_50_ (100 μl volume) of NY-ESO-1 S-FLU (PR8) virus used under inhalation isoflurane anaesthesia. In some experiments, intramuscular infection with 1×10^9^ PFU of Hu Ad5 NY-ESO-1 virus or 5×10^7^ PFU of Fowl Pox NY-ESO-1 virus was also performed.

### Lung cells preparation and CD8^+^T cells enrichment

Single cell preparation from lungs was prepared as described before ^52^. Briefly, mice lungs were perfused with 10 ml of PBS and digested in 0.5 mg/ml of Collagenase type IV in HBSS/10%FBS for 45 minutes after chopping finely with scissors. After digestion, lung tissues were passed through a 19G needle a few times and filtered through a 70 μm cell strainer. After two washes in FACS buffer (PBS containing 1% FBS and 2mM EDTA), the cells were subjected to RBC lysis (Qiagen-RBC lysis buffer) for 5-7 minutes followed by two washes with FACS buffer. For adoptive transfer experiments, naïve CD8^+^ T cells from 1G4 mice spleen were enriched using MACS beads (Pan T cell isolation kit II and CD8a isolation kit, Miltenyi Biotec) and labeled with 5μM CellTrace^TM^ Violet (CTV) (ThermoFisher Scientific) following manufacturer’s instructions. Approximately 2×10^6^ CD8^+^ T cells in 200 μl volume were adoptively transferred to HHD mice.

### *Ex vivo* peptide restimulation assay, intracellular staining and tetramer staining

Splenocytes or lung cells (2 × 10^6^) were isolated from either naïve or infected or tumour-bearing HHD or BALB/c mice and were cultured in the presence of HLA-A2 restricted IAV M1 peptide _158-66_(GILGFVFTL)(Cambridge peptide) or NY-ESO-1 peptide _157-65_ (SLLMWITQC)(Sigma peptide) or H-2K^d^ binding IAV NP peptide _147-55_(TYQRTRALV) (Cambridge peptide) or H-2D^d^ restricted NY-ESO-1 peptide _81-88_ (RGPESRLL) (Cambridge peptide) in complete RPMI-1640 medium supplemented with 10% FBS, 2.1 mmol/L ultra-glutamine in the presence of Brefeldin A (5 μg/ml) and monensin (2 μM; BioLegend). After 5 hours of incubation, cells were stained for extracellular markers (CD3e, CD8a, B220 and CD44) and viability dye ( Near IR dead cell staining kit, Invitrogen. Cells were then stained for intracellular IFNγ, TNFα and IL2 using an Intracellular Fixation and Permeabilization Buffer Set (eBioscience) following manufacturer’s instructions. In some experiments, lung cells or splenocytes were prepared as described above and stained with appropriate surface antigens followed by PE labelled H-2D^d^ restricted NY-ESO-1 peptide _81-88_ (RGPESRLL) and brilliant violet 421 labelled HLA-A2 restricted IAV M1 peptide _158-66_(GILGFVFTL) or H-2K^d^ binding IAV NP peptide _147-55_(TYQRTRALV) (kindly provided by NIH tetramer facility at Emory University). Samples were acquired on a FACScanto II or LSR Fortessa-X50 flow cytometer (BD Biosciences) and data were analyzed with FlowJo version 10.4.1. Compensation beads (eBioscience) were used to generate the compensation matrix, and FMOs were used as control.

### Tumour model

For tumour model, 4T1 metastatic breast carcinoma – syngeneic mouse model has been used as described before ^53, 54^ with some modifications. For prophylactic lung metastasis model, BALB/c mice were intranasally infected with 1×10^6^ TCID_50_ of GFP S-FLU or NY-ESO-1 S-FLU on day 0 followed by booster infection on day 14. Mice were challenged with tail vein injection of 2×10^5^ 4T1-NY-ESO-1cells on day 16 and on day 30 ( day 14 post injection of tumour cells) mice were euthanized and lungs were fixed in Bouin’s solution to count the tumour nodules. For therapeutic tumour model, mice were first injected with 2×10^5^ 4T1-NY-ESO-1 cells on day 0 followed by intranasal infection with GFP S-FLU or NY-ESO-1 S-FLU virus on day 2 and day 10 in the doses mentioned above. Mice were closely monitored and were euthanized on day 16 post tumour cells injection. For the spontaneous metastasis model, 4T1 cells were subcutaneously injected in the flank of BALB/c mice and the mice were intranasally or intramuscularly infected with GFP S-FLU or NY-ESO-1 S-FLU virus on day 4 and 18 post tumour cells injection. In some experiments, immunized mice were also received i.p injection of anti PD-1 antibody (12.5 mg/kg, RMP1-14, BioXCell) or isotype antibody. Tumour volumes were measured every other day until day 28-30 at humane end point (when tumour size reaches >1.2cm^3^). Then mice were euthanized and metastasis colony formation assay (clonogenic assay) was performed to quantify 4T1-derived cells in the lungs of transplanted mice. Entire lungs were minced and incubated in HBSS media with 10% FBS, supplemented with collagenase (1 mg/ml; Sigma-Aldrich,) and deoxyribonuclease (100 μg/ml; Sigma-Aldrich) for 45 min. Thereafter, lung fragments were homogenized through a 100 μm filter and rinsed with 5 ml of PBS. After centrifugation, cell pellets were subjected to RBC lysis (Qiagen RBC lysis buffer) and after 2 times washing in PBS, cells were resuspended in 10 ml of selection media (RPMI 1640 media with 10% FBS and 60 μM 6-Thioguanine (Sigma-Aldrich)) before being diluted 1:10 and 1:100 in selection medium. Cells were plated in culture dishes and the colonies were stained after 14 days with 1% crystal violet/70% methanol solution and counted.

### Statistical Analysis

Unpaired, two-tailed Student’s test, One way ANOVA with Tukey’s multiple comparisons were used to calculate statistical significance (Prism, GraphPad)

## Declarations: Acknowledgements

This work was supported by CRUK, UK grant. The funders had no role in study design, data collection and analysis, decision to publish, or preparation of the manuscript. This paper is dedicated to our wonderful mentor, Prof. Vincenzo (Enzo) Cerundolo.

---

Supplementary Figure 1. Expression of PR8-NP and NY-ESO-1 proteins in NY-ESO-1 S-FLU virus infected HEK cells. HEK cells were infected with NY-ESO-1 S-FLU virus and the cells were fixed and permeabilised after 18hrs of infection followed by staining with DAPI and immune stained for NP and NY-ESO-1 proteins.

Supplementary Figure 2. Intranasally administered NY-ESO-1 S-FLU virus mainly infect lung epithelial cells. (A) Representative FACS plot showing the infected lung cell subsets on day 2 post infection. BALB/c mice were intranasally infected with NY-ESO-1 S-FLU virus and lung cell subsets were analyzed for virus infection by PR8-NP staining. (B) Bar graph (left) showing the frequency of NP^+^ non-immune cells (CD45^-^) and immune cells (CD45^+^) in lungs following infection with NY-ESO-1 S-FLU virus on day 2 and day 4. Bar graph (right) showing the frequency of NP^+^ lung epithelial cells on day 2 and day 4 post infection. Data in Supplementary Figure 2 is representative of two independent experiments with similar results. *P<0.05, **P<0.01(2 -tailed student-t test). Error bars: mean ± SEM

Supplementary Figure 3. *In vivo* T cell priming at draining lymph node following infection with NY-ESO-1 S-FLU virus. (A) Experimental design. n=5-6 mice per group. Data are representative of at least 2 independent experiments with similar results. (B) Histograms represent the analysis of cell trace violet labelled 1G4 CD8^+^T cells proliferation at mediastinal lymph node 3 days post infection with GFP S-FLU or NY-ESO-1 S-FLU virus. The graph on the right shows the respective proliferation index for the proliferating 1G4 CD8^+^T cells. *P<0.05, (2 -tailed student-t test). Error bars: mean ± SEM

Supplementary Figure 4.Intranasal infection with NY-ESO-1 S-FLU virus elicits specific CTL response in spleen. (A) Representative FACS plots showing IFNγ secreting CD8+T cells in spleen. On day 10 post intranasal infection, splenocytes were stimulated *ex vivo* with IAV NP peptide or NY-ESO-1 CTL peptide and IFNγ secreting CD8+T cells were analyzed. (B) Bar graph showing the frequencies of IFNγ secreting CD8+T cells in spleen following *ex vivo* stimulation IAV NP peptide or NY-ESO-1 CTL peptide. The results shown are a representative of three independent experiments with similar results (n = 4-5 mice/group) ***P<0.001, ns- not significant (2 -tailed student-t test). Error bars: mean ± SEM

Supplementary Figure 5 Absolute number of NY-ESO-1 specific CD8+T cells in Lungs and spleen following intranasal or intramuscular infection. (A) Bar chart show the absolute number of IFN γ producing CD8+T cells in lungs in *ex vivo* stimulation with Flu M1 peptide ( left panel) or NY-ESO-1 peptide (right panel). (B) Bar chart show the absolute number of IFN γ producing CD8+T cells in spleen in *ex vivo* stimulation with Flu M1 peptide ( left panel) or NY-ESO-1 peptide (right panel). HHD mice were intranasally or intramuscularly infected with NY-ESO-1 S-FLU virus and Flu M1 specific or NY-ESO-1 specific IFN γ producing CD8+T cells in lungs (left) and spleen (right) were analyzed by *ex vivo* peptide stimulation on day 10 post infection. *P<0.05, **P<0.01 (2 -tailed student-t test). Error bars: mean ± SEM

Supplementary Figure 6 Gating strategy for flow cytometric analysis of lung tissue resident memory T cells.

Supplementary Figure 7 Infection with HA switched NY-ESO-1 S-FLU virus elicited a stronger NP specific CTL responses in spleen A-left) Representative FACS plot showing IFNγ and TNFα secreting and (B-left) NP tetramer positive CD8^+^T cells in spleen. Bar graph show the quantification of IFNγ secreting CD8^+^T cells (A-right) and NP tetramer positive CD8^+^T cells (B-right) in spleen. Balb/c mice were intranasally infected with X31 virus and re-infected with NY-ESO-1 S-FLU (X31) or NY-ESO-1 S-FLU (PR8) on day 24 p.i and NY-ESO-1 and NP specific CD8^+^T cell responses in spleen were analyzed on day 7 post-secondary infection by *ex vivo* stimulation with NY-ESO-1 CTL peptide _81-88_ (RGPESRLL) and NP tetramer staining respectively. Data shown in (A) and (B) are pooled results of two (n=8 mice/group) and three independent experiments (n = 12 mice/group) respectively. ****P<0.0001 and ns- not significant (one-way ANOVA-multiple comparisons). Error bars: mean ± SEM.

Supplementary Figure 8 T cell responses elicited by intramuscular injection of different virus vaccines. Bar graphs show the percentage of IFNγ secreting CD8^+^T cells in lungs (left) and spleen (right) in *ex vivo* stimulation of lung cells or splenocytes with NY-ESO-1 CTL peptide _81-88_ (RGPESRLL). Balb/c mice were intramuscularly infected with 1×10^7^ TCID_50_ NY-ESO-1 S-FLU virus or 1×10^9^ PFU Hu Ad5 NY-ESO-1 virus or 5×10^7^ PFU Fowl Pox NY-ESO-1 virus and NY-ESO-1 specific CTL responses were analyzed in lungs and spleen on day 10 post infection. The results shown are representative of two independent experiments with similar results (n = 5-6 mice/group). ** P<0.01, and ***P<0.001 and ns- not significant (one-way ANOVA- multiple comparisons). Error bars: mean ± SEM.

