## Supplementary material for "Recombinant single-cycle influenza virus as a new tool to augment antitumour immunity with immune checkpoint inhibitors": Suppl.Figures

**Supplementary Figure 1.**
**Expression of NY-ESO-1 protein in NY-ESO-1 S-FLU virus infected cells**

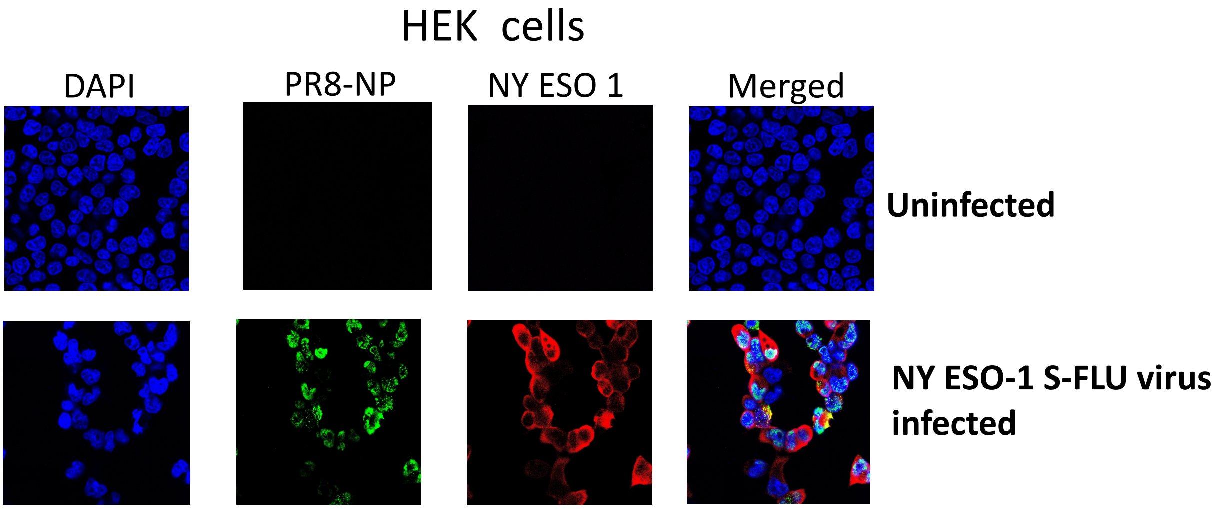

#### Supplementary Figure 2

**Intranasally administered NY-ESO-1 S- FLU virus mainly infect lung epithelial cells**

A)

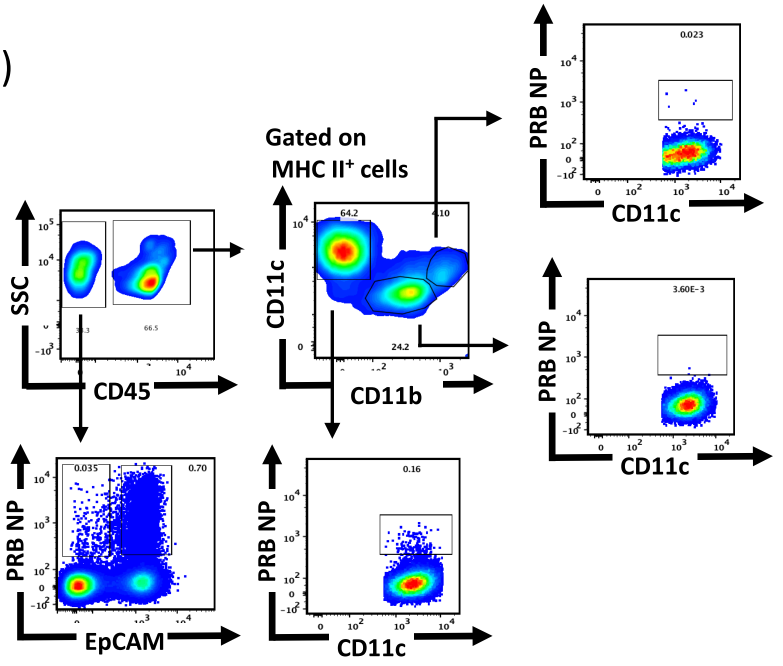


B)

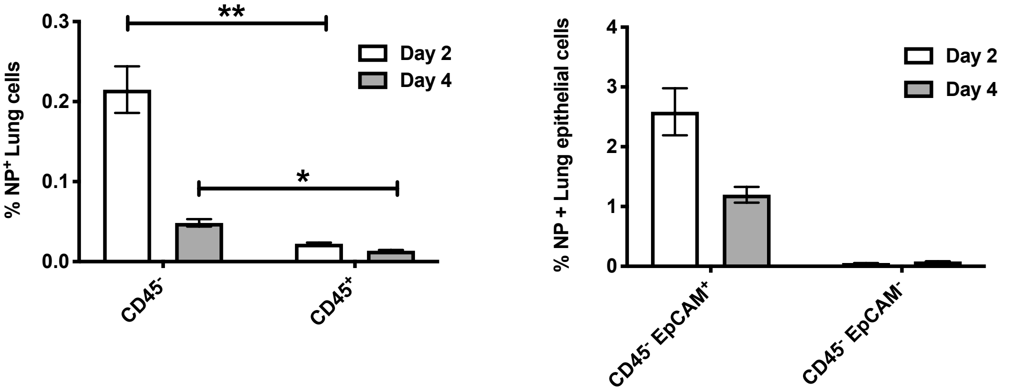

### Supplementary Figure 3

#### *In vivo* T cell activation following infection with NY-ESO-1 S-FLU virus

A)

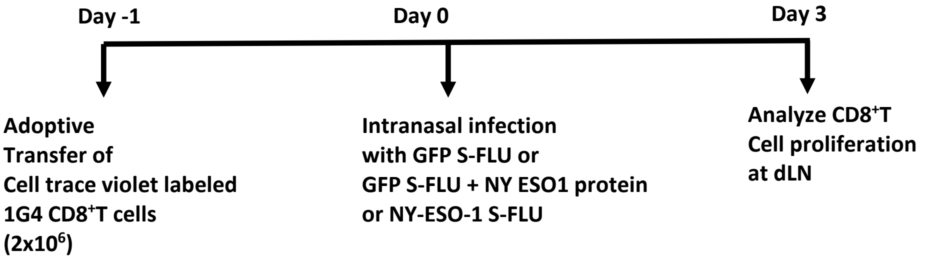


B)

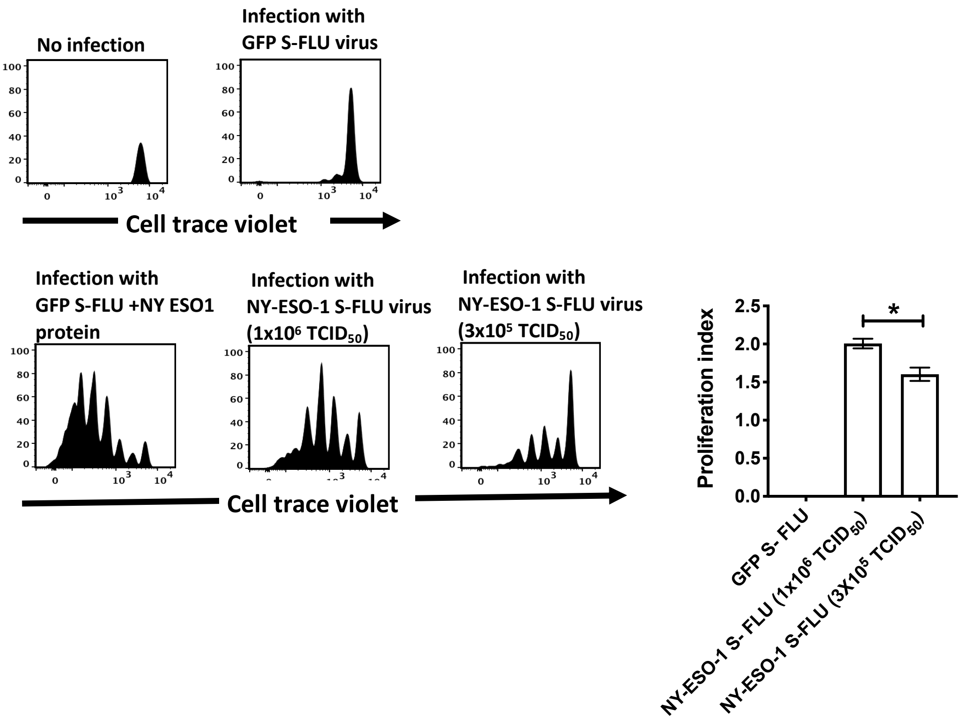

### Supplementary Figure 4

#### Intranasal infection with S-NY- ESO 1 FLU virus elicits specific T cell response in spleen

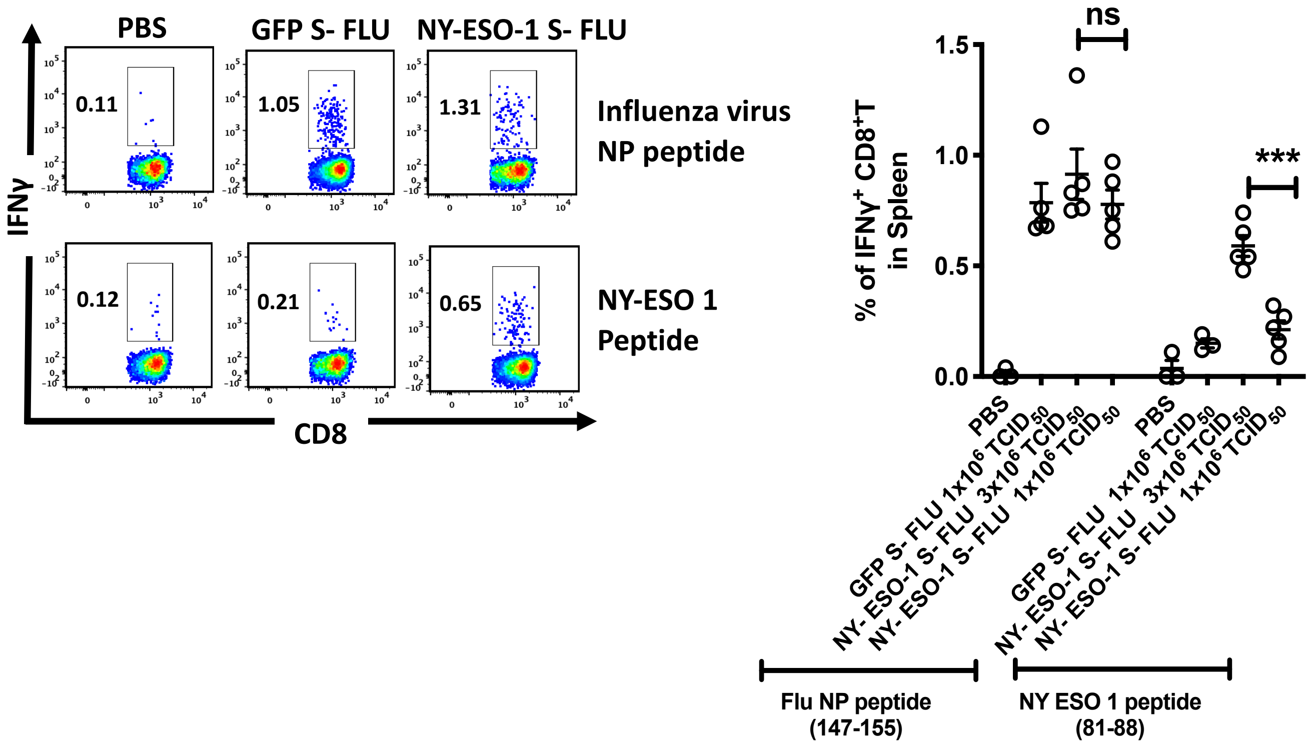

#### Supplementary Figure 5
#### Absolute number of NY-ESO-1 specific CD8+T cells in Lungs and spleen following intranasal or intramuscular infection

A)

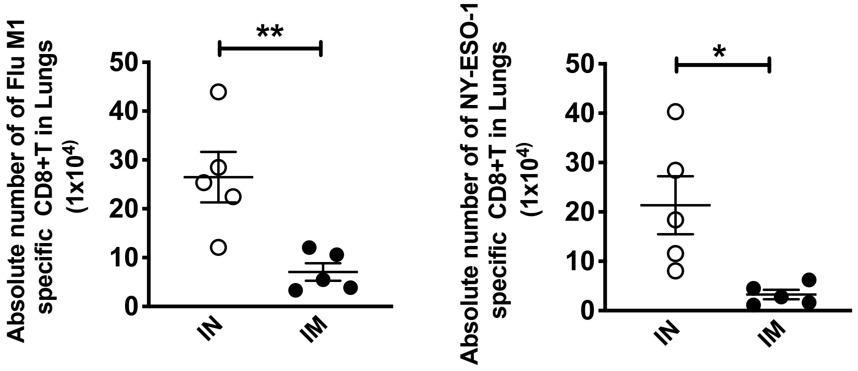


B)

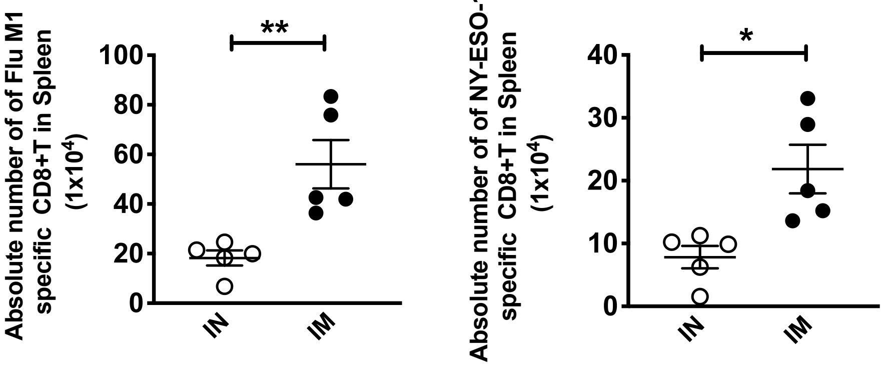

### Supplementary Figure 6

#### Gating strategy for flow cytometric analysis of lung tissue resident memory T cells

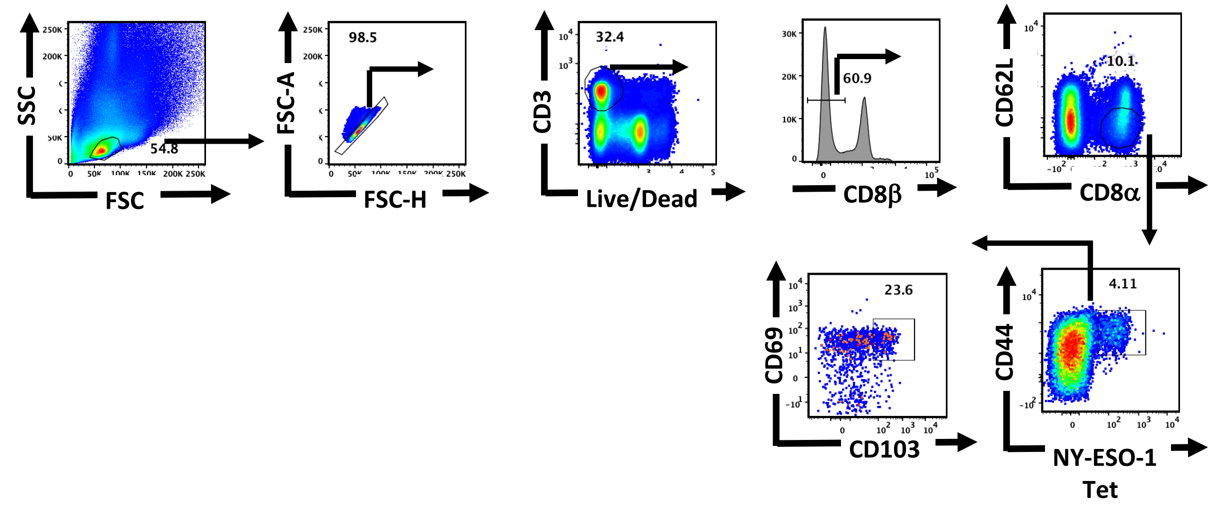

**Supplementary Figure 7**

**Infection with HA switched NY-ESO-1 S-FLU virus elicited a stronger NP specific CTL responses in spleen**

A)

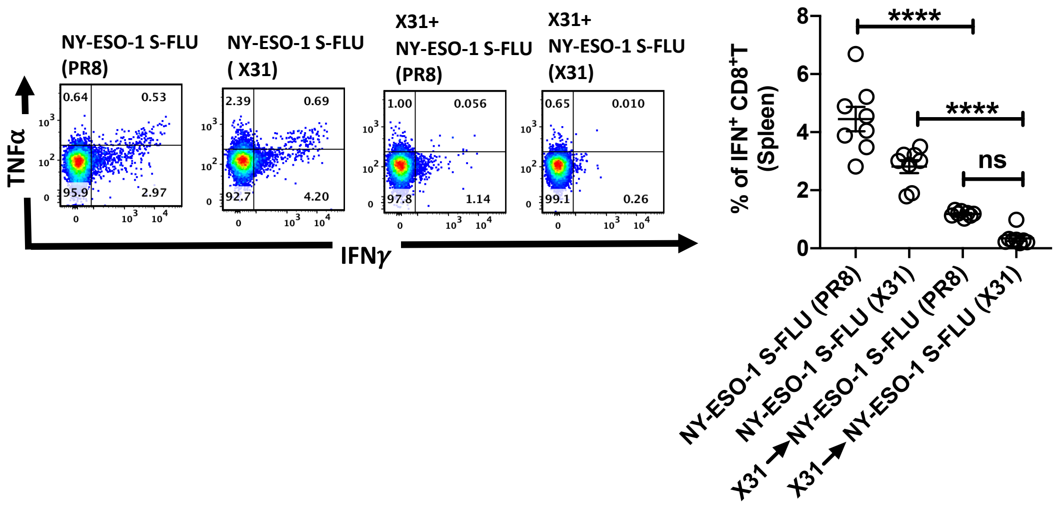


B)

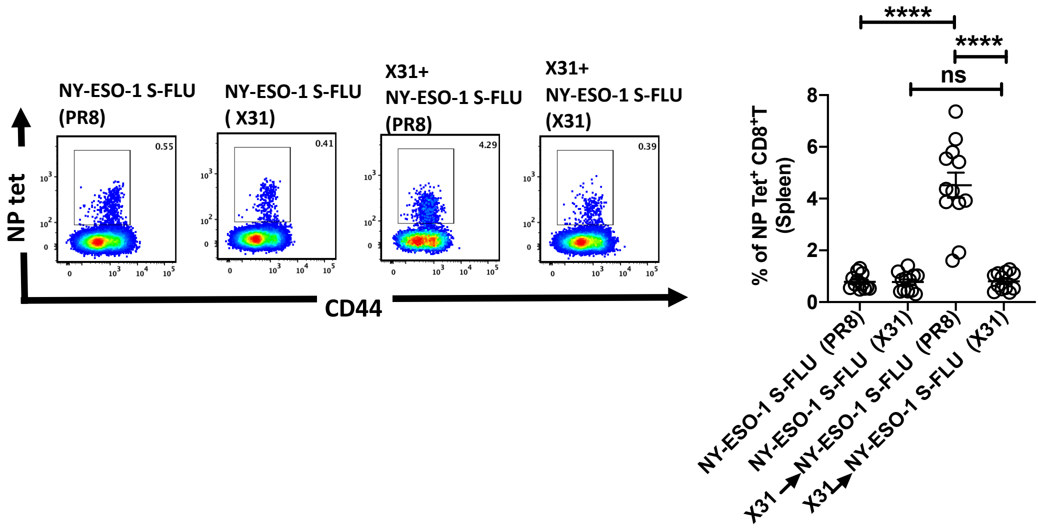

#### Supplementary Figure 8
#### T cell responses elicited by intramuscular injection of different virus vaccines

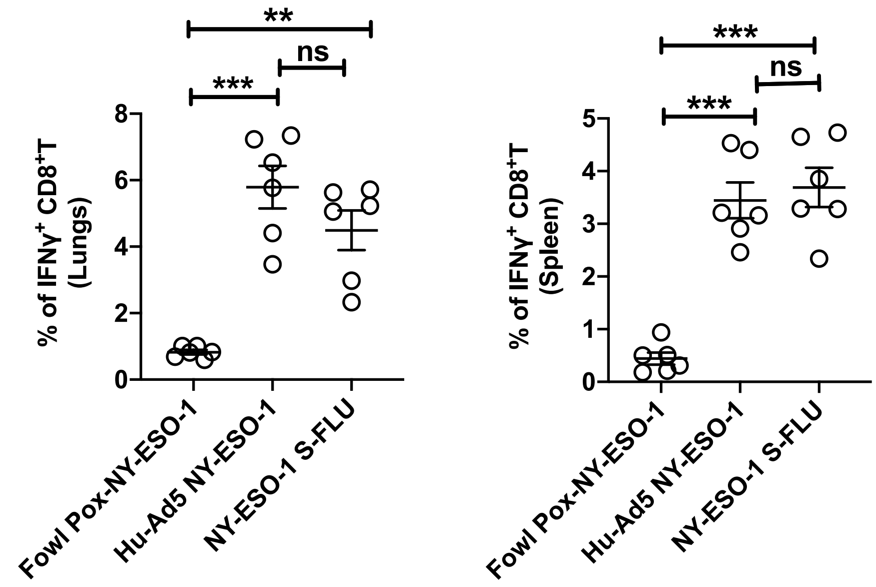
